# Genome-Wide Architecture of Adaptation in Experimentally Evolved *Drosophila*

**DOI:** 10.1101/2020.10.30.361857

**Authors:** Thomas T. Barter, Zachary S. Greenspan, Mark A. Phillips, José M. Ranz, Michael R. Rose, Laurence D. Mueller

**Affiliations:** Department of Ecology and Evolutionary Biology, University of California, Irvine, Irvine, CA 92697; Department of Integrative Biology, Oregon State University, Corvallis, OR 97331

## Abstract

The molecular basis of adaptation remains elusive even with the current ease of sequencing the genome and transcriptome. We used experimentally evolved populations of *Drosophila* in conjunction with statistical learning tools to explore interactions between the genome, the transcriptome, and phenotypes. Our results indicate that transcriptomic measures from adult samples can predict phenotypic characters at many adult ages. Importantly, when comparing the genome and transcriptome in predicting phenotypic characters, we find that the two types of data are comparably useful. When using genome sites as predictors for the expression of the transcriptome, we find that gene expression is influenced by genomic regions across all large chromosome arms. Conversely, we found many genomic regions influencing the expression of numerous genes, which is consistent with widespread pleiotropy. Our results also highlight the power of the combination of experimental evolution, next-generation sequencing, and statistical learning tools in exploring the molecular basis of adaptation.

## Introduction

Despite recent attempts (1, 2), the study of the molecular architecture of adaptation is still in its infancy (3). Experimental evolution establishes selection on populations in a well-defined environment as a means to decipher the interplay between population genetics and selection (4, 5, 6). Genome-wide sequencing can provide extensive catalogs of changes in the frequency spectrum of genetic variants in lab evolved populations, as well as a portrait of their expression levels (7, 8). New statistical approaches have been developed that capitalize on genome-wide sequencing in experimental evolution (9, 10, 11). The hope is that the combination of high-throughput sequencing, statistical learning, and well-characterized life-history characters might help us to understand how genetic variation underpins adaptation generally.

Here, we study the interplay between genomics, transcriptomics, and life history traits from 20 experimentally evolved *Drosophila melanogaster* populations. Of these populations, ten populations have been selected for accelerated development, and the remaining ten have been selected for postponed reproduction (Fig. 1). For all populations, we previously sequenced their genomes (1) and female transcriptomes (12) and collected data on their life history traits (13). In this study, we use genomic and transcriptomic data to infer which gene regions might be causally linked to the phenotypes of age-specific mortality and fecundity, the life-history traits that numerically define biological fitness. In a similar vein, we use genomic data to infer which loci are causally linked to changes in gene expression across the transcriptome. These inferences are made using a statistical learning technique called fused lasso additive model, or FLAM (10), which theoretical work suggests is well suited to the task (11).

**Fig. 1.**
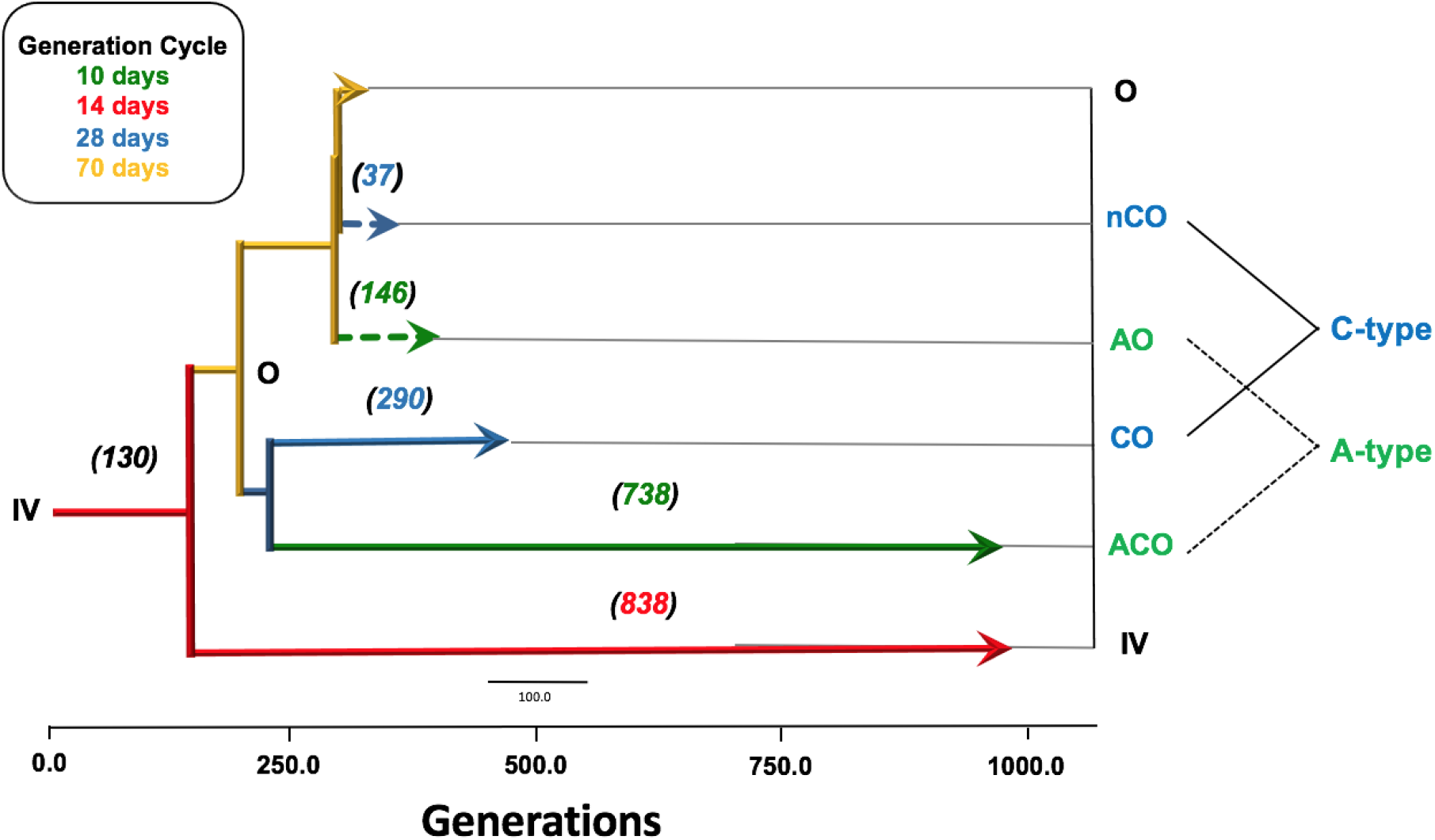
Twenty population selection history. All treatments in the study share ancestry. Each color represents a different selection regime and population type (A and C). A-type populations have a 10-day life cycle, where the C-type populations have a 28-day life cycle.

## Results

Using phenotypic, genomic, and transcriptomic data (Fig. 2 and 3; Material and Methods), we investigated the potential interactions among these types of data.

**Fig. 2.**
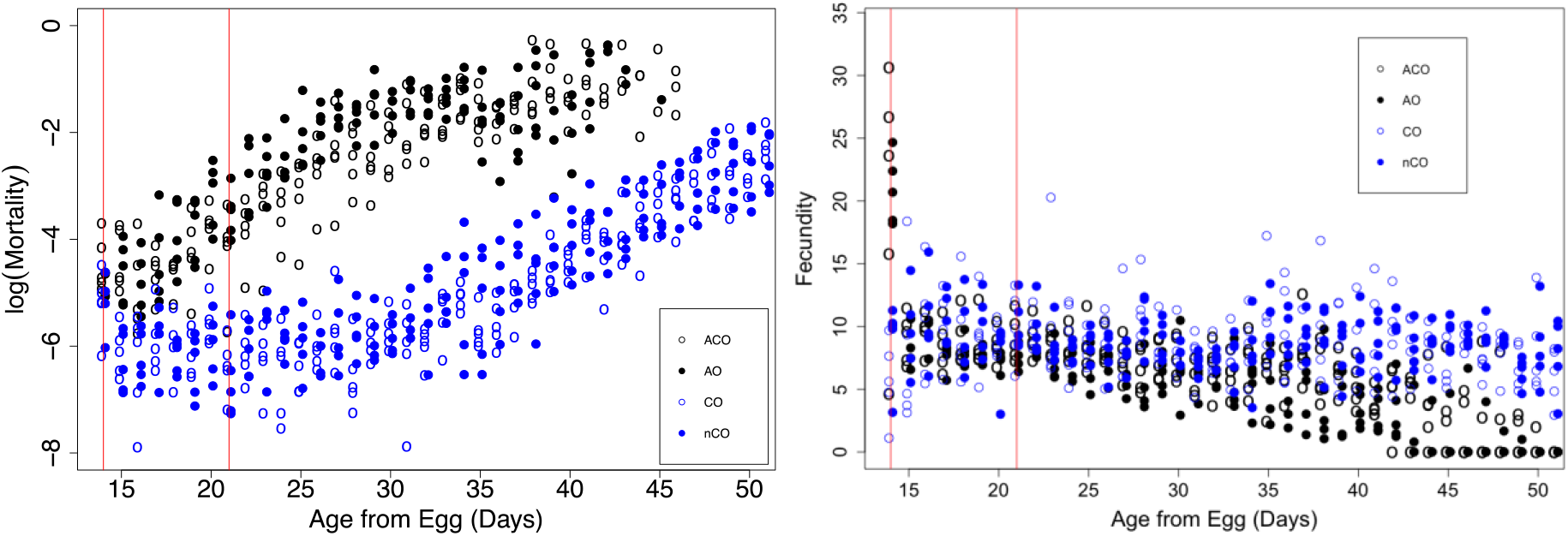
Age specific phenotypes for twenty large cohorts, one for each of twenty populations. Female age specific mortality (Burke et al. 2016) is shown on the left and the age specific fecundity (Burke et al. 2016) is shown on the right. Black dots signify A-type populations and blue dots represent C-type populations. Open circles represent long-standing populations and closed circles represent recently-derived populations. The red lines represent the time of sequencing for the transcriptomic work.

**Fig. 3.**
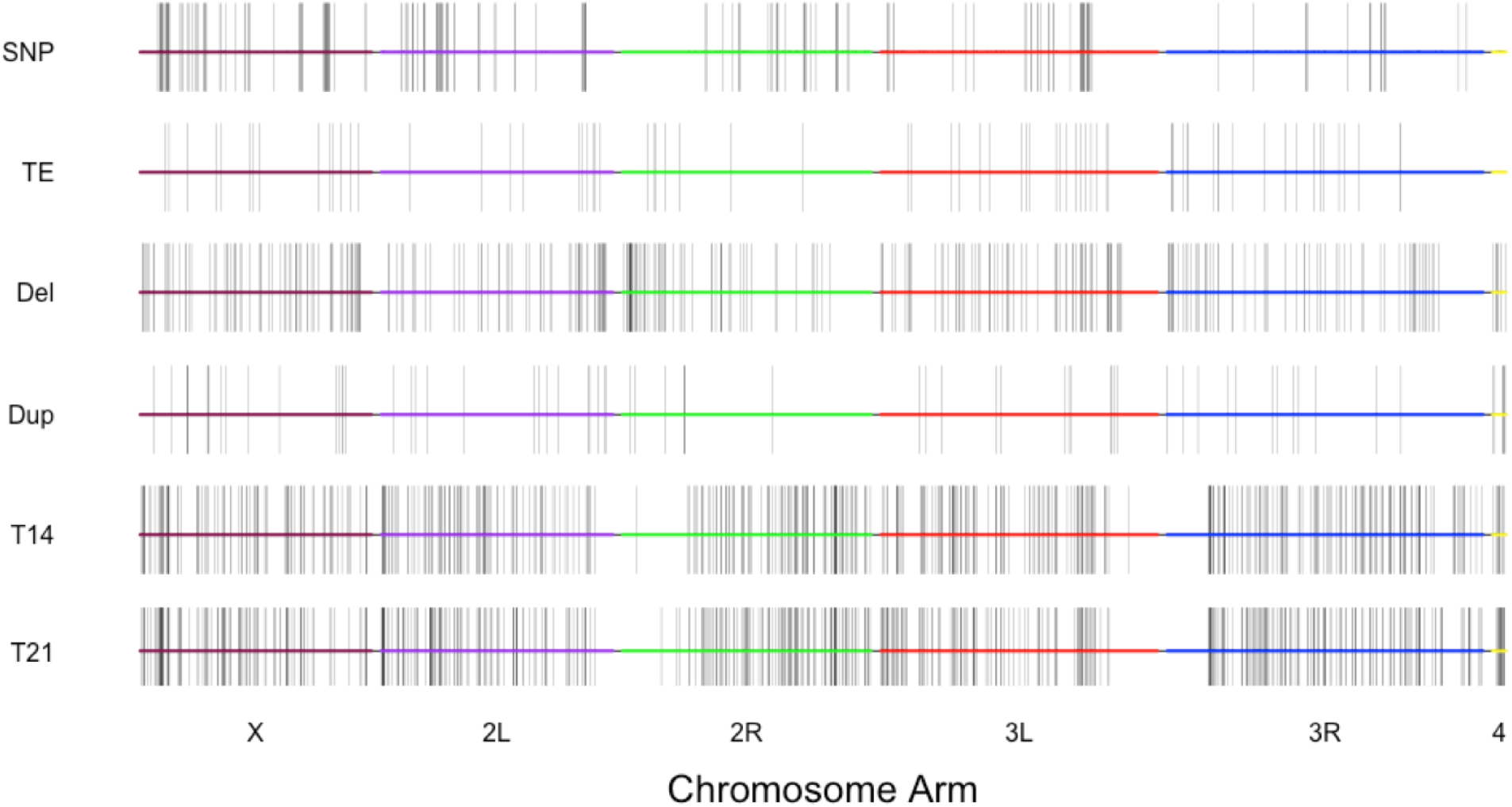
Chromosomal distribution of the molecular differences between the two treatments. Each bar shows the position of differentiation across the genome and transcriptome. From the top to the bottom each panel shows differentiation in SNPs, TEs, deletions, duplications, day 14 expression, and day 21 expression respectively.

### Predicting age-specific mortality and fecundity using genomics and transcriptomics

It is conceivable that both the genome and the transcriptome have the ability to predict a given phenotype, but here we ask whether the genome or transcriptome has more predictive power. Using FLAM, we evaluated whether genomic and transcriptomic data performed better in providing viable predictors, which might suggest which is more relevant for understanding the molecular basis of adaptation. To address this, and with the purpose of predicting mortality and fecundity at days 14 and 21 from egg, we performed three different analyses in which we used different types and combinations of data as predictors: (i) genomic data (SNP frequencies); (ii) normalized gene expression levels; and (iii) both. The results varied depending on which phenotype was used as outcome (Fig. 4). For day 14, it is evident that the genomic data appears to be better at predicting the age-specific mortality data, but the transcriptomic data appears to be better at predicting the age-specific fecundity data (Fig. 4). For the most part, it appears that the genomic locations and the genes found as optimal predictors of phenotypes individually also show up in the combined list. Neither genomic nor transcriptomic data are obviously better at predicting day 14 phenotypes.

**Fig. 4.**
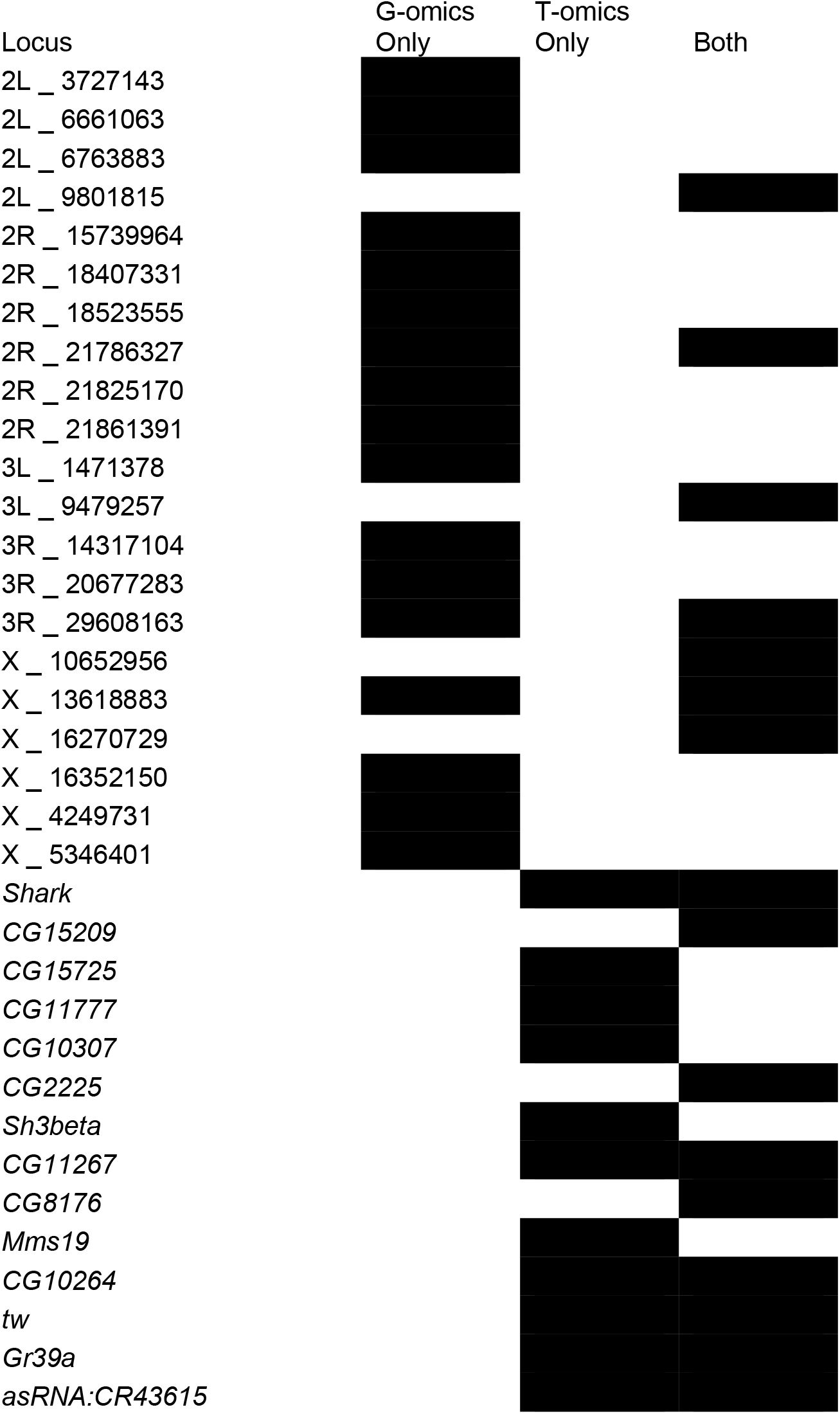
Genomics versus Transcriptomics in predicting mortality at day 21. Predictive loci (black box) from genomics (G-omics) and transcriptomics (T-omics) in all three analyses and from which analyses they are considered predictive loci. For SNP genomic regions, the molecular coordinate shown corresponds to their middle point in the form of chromosome arm and position. These genomic regions are classified as 50kb windows which contained at least 3 differentiated SNPs between the population types A and C.

As no statistically significant differences in fecundity were found at ages greater than 21 days (13), day 21 genomic and transcriptomic data were used to predict mortality only. We found that both – omic data sets performed relatively well (Fig. 4). In contrast to day 14, the model does not appear to strongly favor one −omic data set over the other. As for days 14 and 21 we found that both genomic and transcriptomic data harbor optimal predictors, it seems ill-advised to use only one type of −omic data. The reason why one type of −omic data may perform better for some phenotypes but not for others remains is not apparent at this time.

### Transcriptome plasticity

Unlike the genome, the transcriptome is subject to change over the course of an organism’s life, most notably during developmental transitions. This is the case for *D. melanogaster*, but once flies reach adulthood the transcriptome shifts to a more static state, similar to that of the genome (14), due to minimal cellular changes during a fly’s adulthood. [Few cells divide in the adult soma, and there is relatively little protein synthesis (15).] If the adult transcriptome is static, then assaying multiple time-point transcriptomes for adult *Drosophila* may be redundant.

We used the expression levels of differentially expressed genes from day 14 and day 21 as predictors of mortality at days 14-35, while we used the expression levels of differentially expressed genes from day 14 as predictors of fecundity at days 14-35. From our analyses at day 14 and day 21, FLAM reported which genes were classified as viable predictors and these predictors were then labeled as focal predictors. Our data was then fractured into two parts: 16 populations were used as a training set and the remaining 4 populations were used as a testing set. The splitting of populations for a training set and testing set was repeated for different combinations of populations to ensure each population was included in the testing set. Using our training sets of populations, we include all age-specific mortality and fecundity data as outcomes and the expression values for the focal predictors as inputs. By doing this, we restricted FLAM to using only these focal predictors determined from our analyses at day 14 or day 21. With each specific fit, we use expression levels from the test set to predict their phenotypes and compare it to the observed phenotypic values. The correlation between the predicted phenotype and observed phenotype was then determined (Figure 5).

**Fig. 5.**
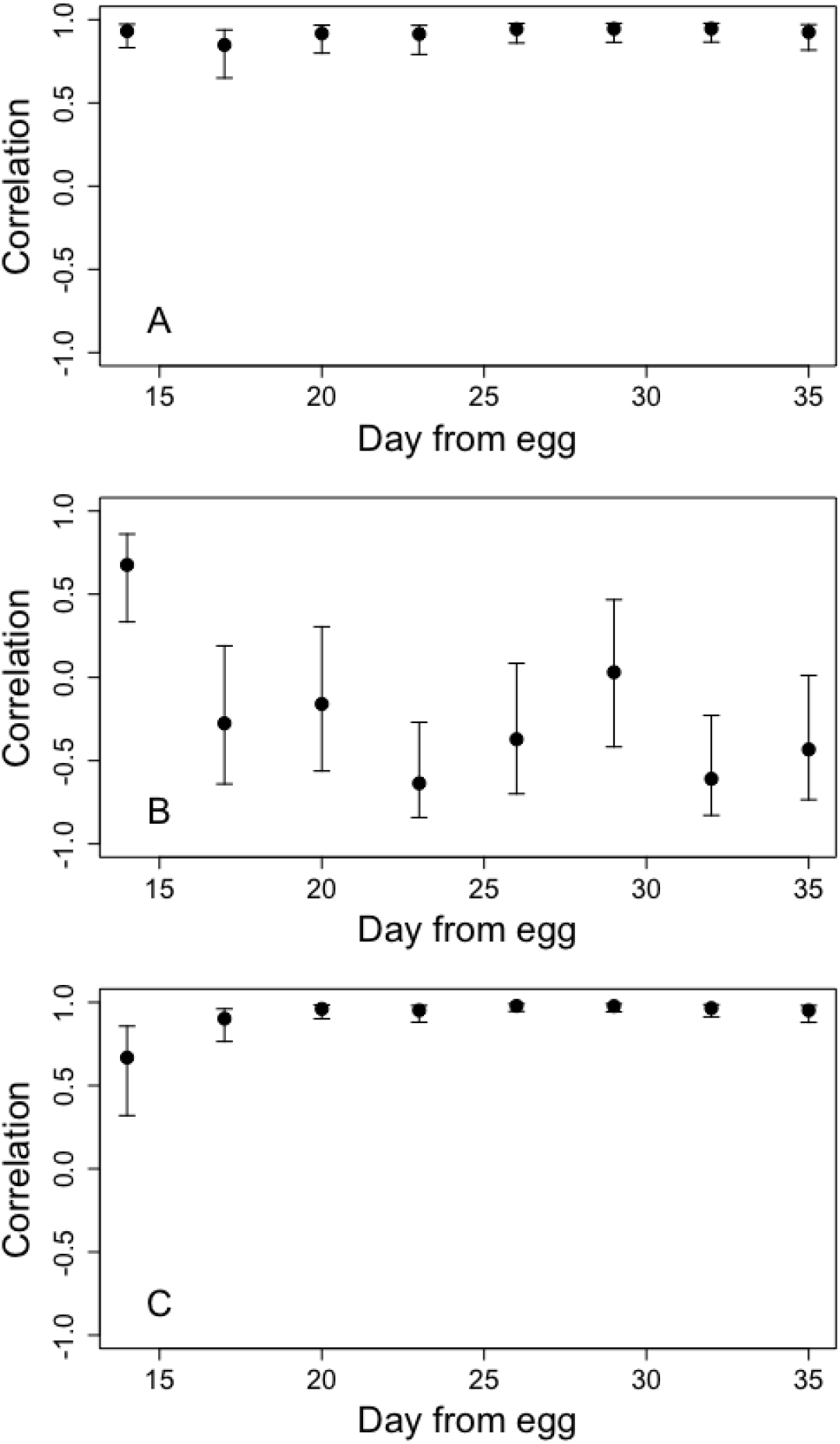
Correlation values when comparing predicted phenotypic values to true phenotypic values at every age. We compare how well focal transcriptomic predictors can predict phenotypes at non-focal ages. The values shown correspond to the comparison between the actual phenotypic values and the predicted phenotypic values from our test set. The predicted phenotypic values were generated using the model fit at each age from the focal transcriptomic predictors. (A) Day 14 transcriptomic data were used as predictors and age-specific mortality as outcomes. (B) Day 14 transcriptomic data were used as predictors and age-specific fecundity as outcomes. (C) Day 21 transcriptomic data were used as predictors and age-specific mortality as outcomes.

We find that the transcriptomic data reasonably predicted mortality at all ages, even though the transcriptomic data were limited to the focal genes identified from either day 14 or day 21. However, day 21 transcriptomic data appear to be able to predict latter age-specific mortality marginally better. This marginal increase in predictability may originate from the fact that at day 14 the C populations may still have lingering developmental effects, since day 14 is shortly after the C populations have completed developmental transitions and the adults have eclosed.

The same trend is not seen for fecundity. FLAM could only create a good fit for day 14 data. There is no differentiation between the A and C population fecundity values at days 17-25. It appears that there is only a strong correlation between the actual phenotype and the predicted phenotype at the focal age, 14. This is plausible considering the fact that fecundity is significantly different at days 14-16, but not between days 17-25 (Fig. 5). Even though fecundity is significantly different for days after day 25, we still see only a weak correlation between the actual phenotype and the predicted phenotype. This is likely due to the fact that day 14 transcriptomic data do not reasonably predict fecundity at all ages.

### Unbiased approach to predicting transcriptomic expression using genomic variation

Transcription is shaped by both local regulatory sequences (*cis*-) and distantly encoded regulatory factors (*trans*-) (16). Although we do not focus on detailed mechanisms of gene regulation here, we used different types of genetic variation as predictors and each individual transcript’s expression phenotype as outcomes. By doing this, we were able to assess how well SNP regions, transposable element insertions, small indels (<10kb), or duplications of DNA material performed as predictors for the expression of each differentiated transcript. First, we included each type of genomic variant (e.g. SNPs) separately to characterize their effects on the transcriptome. Subsequently, we included all genomic feature variants together to determine which variants were the strongest determinants of mRNA abundance.

In our first analysis, we uncovered numerous causal interactions between the genomic variants and expression levels (Fig. 6, Supplementary Fig. 1-7). These interactions often involved predictor loci located on chromosomes different from that in which the differentially expressed transcript resided. The average number of optimal predictors varied depending on the time point (day 14 and day 21) and on the type of genetic variant (see Supplementary Table 1).

**Fig. 6.**
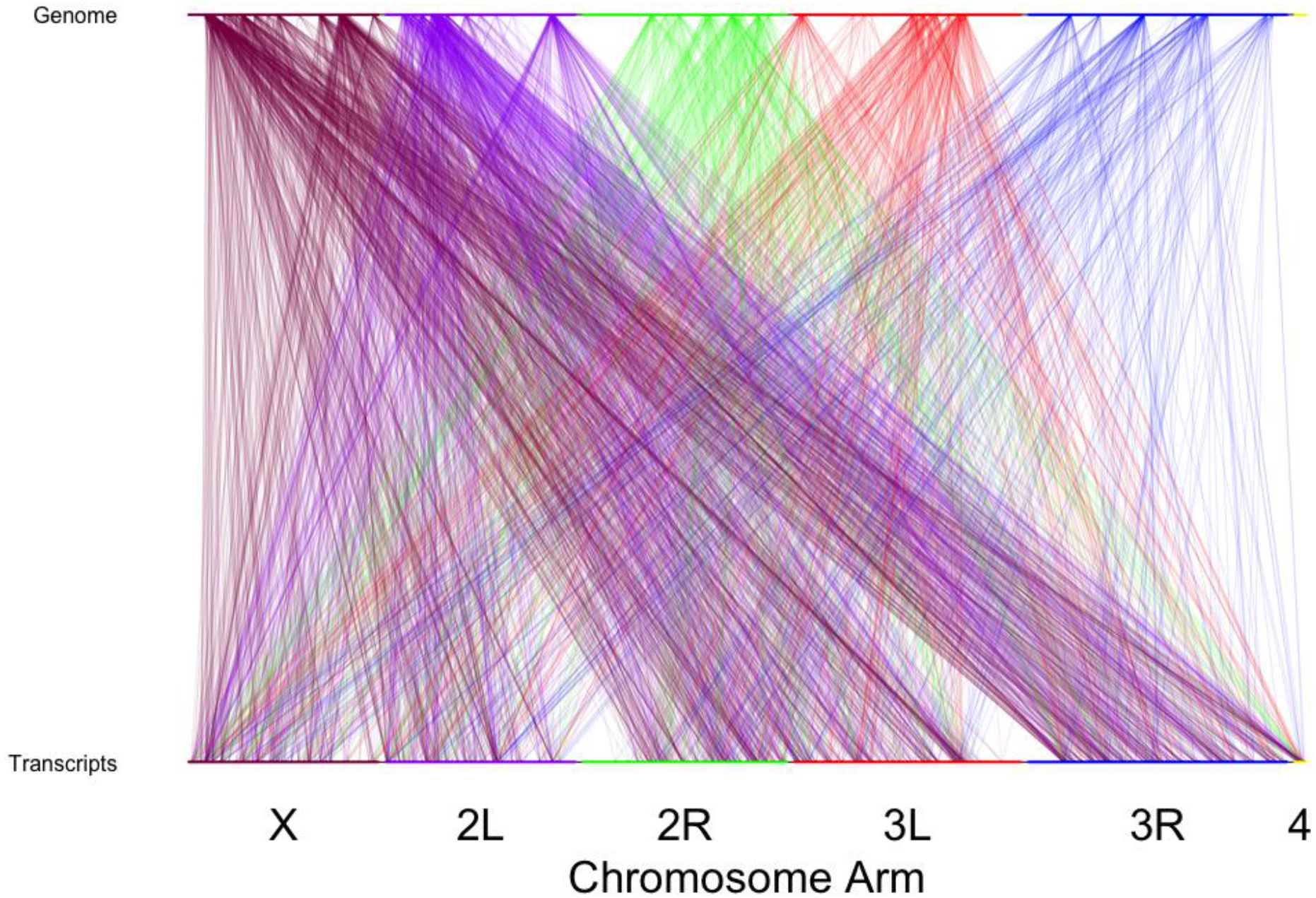
Interactions between predictive SNP regions and differentiated transcripts at day 21 across the *D. melanogaster* chromosomes. Each line maps a predictive SNP region to a transcript for which the SNP region accurately predicts expression. The color of the line denotes which chromosome arm the genomic region originates from. The genomic regions are classified as 50kb windows which contained at least 3 differentiated SNPs. The transcripts are those classified as significantly differentiated for quantitative expression.

Our results show evidence of extensive pleiotropy, in that single differentiated genomic regions are reliably contributing to the prediction of expression in numerous genes. Conversely, we see that the differentiated expression of a single gene is being predicted by numerous differentiated genomic loci, including many that are located on different chromosomes. Although we see many interactions between the genome and the transcriptome, these interactions should not all be considered “regulatory” due to the fact that the genomic regions are quite large, and the mechanistic details of how these regions are potentially affecting gene expression remain unknown. For example, it is conceivable that some predictive genomic regions might have an indirect effect on the differentiated genes between populations, which in turn would result in differential expression.

In our second analysis, we combined the SNP, TE insertion, small indel, and duplication data sets into one large set of potential predictors for differential expression (Fig. 7). We found that the SNP data best predicted the transcriptome differentiation in expression levels between the two types of populations assayed, as FLAM only used the SNP data when all the different data were combined. In other words, the duplication and TE insertion data were ignored by FLAM. A plausible explanation for this result is that TE insertions and duplications were not appreciably variable within each of the two selection treatments. By contrast, the SNP data featured ample variation among the ten populations within each treatment. Given that the expression of the transcripts is variable within treatments, the genomic data that lack variation within treatments will be discarded by the FLAM algorithm as their predictive power is extremely limited without such variation. This is not to say that TE insertions and duplications do not contribute to changes in gene expression, but their lack of differentiation among replicates within these two groups of populations makes them poor predictors of quantitative differences in expression compared to the SNP data.

**Fig. 7.**
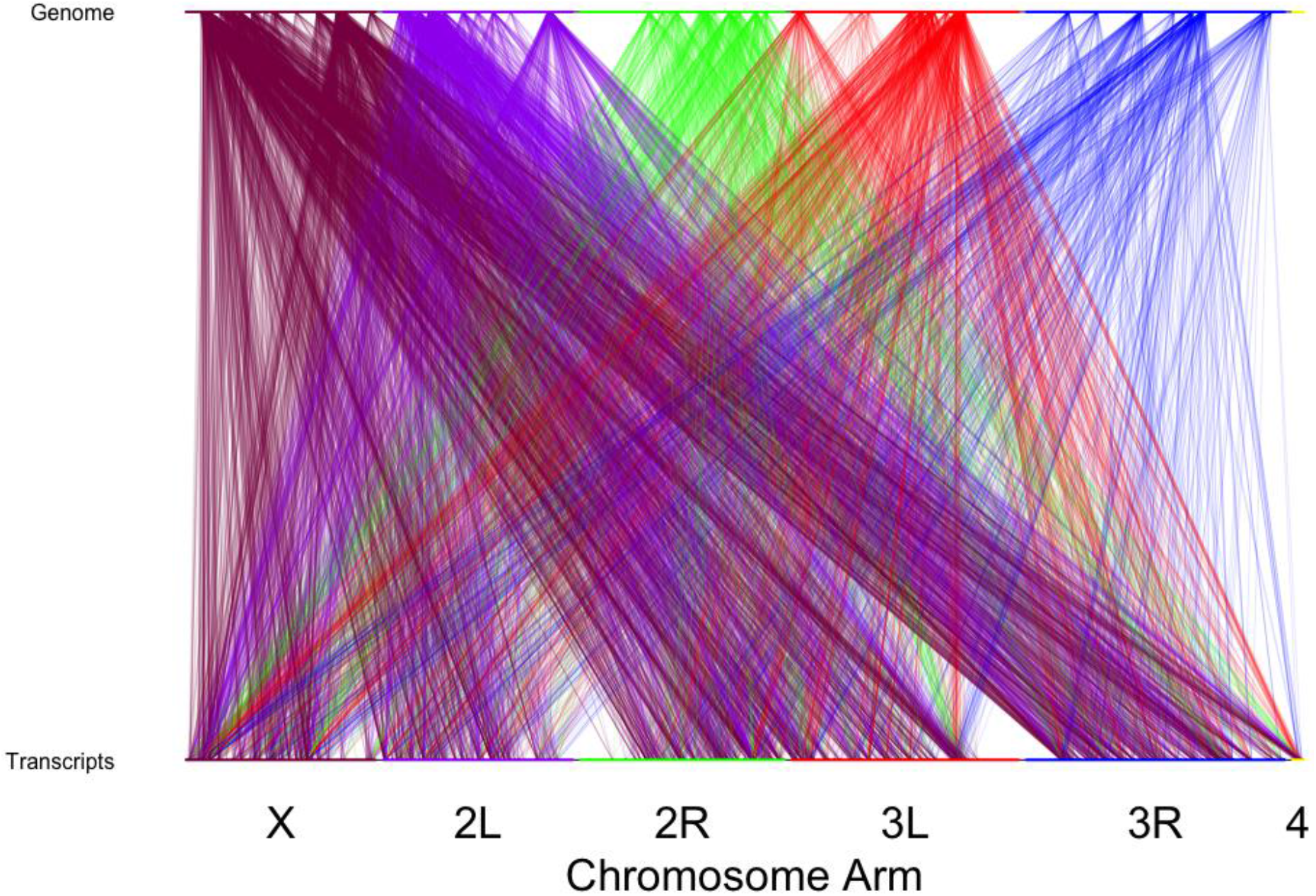
Interactions between predictive genomic character and differentiated transcripts at day 21 across the *D. melanogaster* chromosomes. Each line is an interaction between a predictive genomic character to a gene that the frequency of the genomic character can accurately predict the expression. The color of the line denotes which chromosome arm the genomic character originates from. The genomic characters are classified differently depending on the character. The transcripts are those classified as significantly differentiated for quantitative expression.

### Accuracy of FLAM predictors

For each of the previous analyses, we used each predictor list to determine the accuracy of our FLAM analyses by compiling numerous training and test sets with our genomic and transcriptomic data. We found that all data sets have similar average correlation between the actual expression values and the predicted values from the test sets (Fig. 8, Supplementary Fig. 8-14). The SNP data gave averages of 0.667 and 0.608 correlations between the true expression values and the predicted expression for day 14 and day 21, respectively. The transposable element data gave averages of 0.641 and 0.6134 correlations between the true expression values and the predicted expression for day 14 and day 21, respectively. The insertion and deletion data gave averages of 0.669 and 0.655 correlations for day 14 and day 21, respectively. Lastly, the duplication gave averages of 0.575 and 0.581 correlations for day 14 and day 21, respectively.

**Fig. 8.**
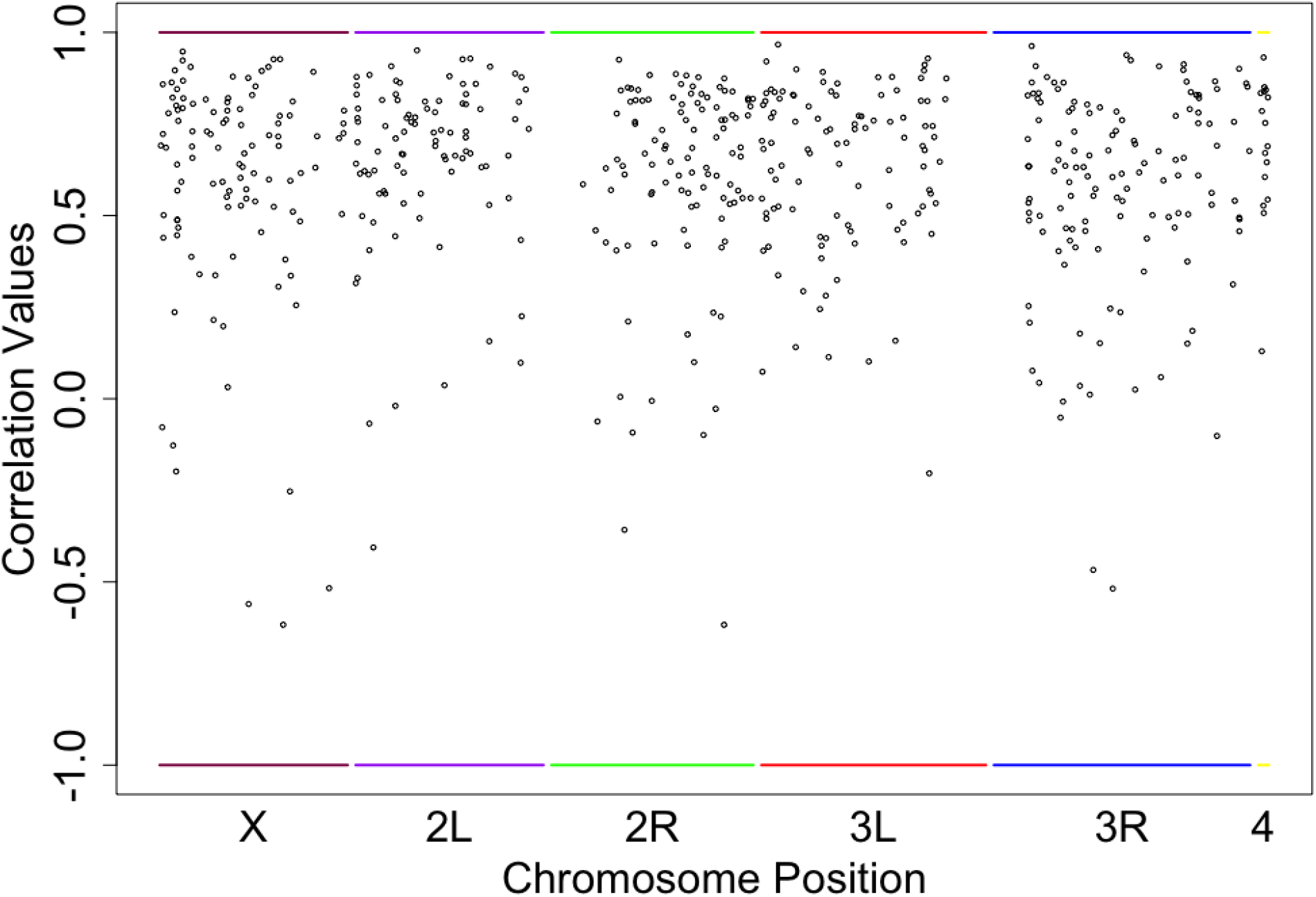
Correlation between predicted value and actual value for each differentially expressed gene. Correlation values were calculated using the predicted expression of each differentiated gene from SNP frequencies and the actual expression values of each differentiated gene at day 21. Each correlation value was plotted at the location of the differentiated gene above. High positive correlation signifies the FLAM model accurately predicted the actual phenotypic value.

Given that the SNP data were preferentially used to predict the expression of the transcriptome when all the genomic data were pooled together, it was unexpected that each type of data is performing at a similar level of accuracy. It is conceivable that the measurement FLAM’s accuracy is masked by the nature of our populations. Our populations are highly differentiated for each transcript that is included in these analyses (12). With this substantial differentiation between groups of populations, it is simple to achieve a moderate correlation between the actual and predicted values using any type of genetic variation, so long as such variation is correlated with treatment. With this in mind, the added predictive power of having within-treatment variation that is found in our SNP data is not truly demonstrated in these tests. Nevertheless, it is evident that the SNP data have superior predictive power over the other data sets from the previous analyses, when all the data sets were included.

## Discussion

Having two clearly defined sets of ten experimentally evolved *D. melanogaster* populations in conjunction with a full suite of genomic, transcriptomic, and phenotypic data for both sets of populations has enabled us to piece together how these three levels of biological machinery interact with one another. Specifically, the 10 A-type populations are clearly differentiated from the 10 C-type populations across the genome (1), across the transcriptome (12), and in age-specific mortality and fecundity (13). Conversely, there is little to no differentiation between the populations within a single set of 10 populations for all three sets of data. This level of differentiation between two sets of populations, together with a high level of convergence within each set, has yet to be seen in sexually reproducing populations in other experiments (17). Lastly, these two sets of populations are closely related, despite their marked differentiation at all three levels, genomic, transcriptomic, and phenotypic.

When reviewing the plasticity of the transcriptome, we found that the transcriptomic data at day 21 accurately predict mortality for all ages after day 21, which is consistent with a relatively quiescent transcriptome once the individuals of a given population type have reached sexual maturity. Since the transcriptome data did not result in accurate predictions of fecundity after day 14, we were unable to make any inferences about the relative stability of transcriptome effects on fecundity at later ages. Further, when comparing the genome and the transcriptome in predicting phenotypic outcomes, we find that there is no clear finding concerning which −omic data set performs better. As it stands, both data sets are used by the machine learning algorithm to predict phenotypic outcomes.

We sought genomic regions that were good predictors for the expression of each differentiated transcript. Although these regions may have predicted the expression of the differentiated gene, these regions should not be considered specific regulators for the gene. Some of these regions may contribute to the regulation of the gene, but it is unlikely that that these regions have evolved solely for their effects on transcript regulation. In addition, the location of each predictive genomic region was not restricted to the *cis*-locale of the gene. It is very clear that gene expression is characteristically affected by many sites across the entirety of the genome. Lastly, it appears as if the X and the 2L chromosome arm (Fig. 6) are favored in the number of predictive genomic regions. However, this may have arisen because those two chromosome arms had the most candidate genomic regions. In other words, the number of predictors from a given chromosome arm is proportional to the number of sites in its sequence that are associated with evolutionary divergence between the “A” and “C” selection regimes.

Currently, we only have the full suite of genomics, transcriptomics, and phenotypic data for 20 populations. As shown in Mueller et al. (11), a 20-population analysis is barely sufficient for detecting causal loci, and by no means will detect the full range of causally important sites in the genome. Ideally, the number of populations used in analyses of this kind should approach 100 populations. Only at such high levels of replication is it plausible that this experimental strategy will reveal a high proportion of the genomic sites that are involved in the response to selection. Although having the full suite of all three types of data is ideal, just having genomics and phenotypic data in additional populations would allow a drastic increase in power for detecting causal loci. With the addition of more experimentally evolved groups of populations we can approach the level of 100 populations, at which point thorough penetration of the genomic complexity of adaptation might be achievable.

## Methods

### Experimental populations

The populations used in this study were subject to two selection regimes which differed with respect to age-at-reproduction (18, 13, 1). Each selection regime was applied to two sets of five populations, each with known distinct evolutionary histories (Fig. 1). The ACO and AO populations, collectively called A-type, were selected for accelerated development and have a generation length of 10 days. The CO and nCO populations, collectively called C-type, have a generation length of 28 days.

### Genomic Data

We used genome-wide SNP, transposable elements, and structural variant data previously published in Graves et al. (1). The details of extraction, sequencing, and read mapping are described in Graves et al. (1). The transposable elements and structural variant data were taken as presented in Graves et al. (1), but the SNP data were realigned with an updated reference genome and statistical tests were rerun. The updated SNP data were then altered slightly to account for linkage disequilibrium to minimize the effects of linkage disequilibrium on our analyses. Briefly, we opted to establish candidate SNP regions around our list of 4,211 candidate SNPs rather than using each candidate SNP. We first divided each chromosome arm into 50 kb windows and discarded any windows that contained less than three candidate SNPs. For the remaining windows, we recorded the position in the window with the smallest *p*-value from the CMH tests of Graves et al. (1). This resulted in a list of 194 positions that serve as representatives of the 50 kb genome regions, *i.e*. 9.7 Mb in total, that met our criteria. These positions and their associated SNP frequencies were then used as inputs in our analyses.

### Transcriptomic Data

We used previously published stranded PE 75 RNA-seq data corresponding to day 14 and day 21 from egg (12). Details about extraction, sequencing, and read mapping are given in Barter et al. (12). After mapping sequencing reads, alignment post-processing was performed with SAMtools v.0.1.19 (19). Read counting per gene and population was done using HTSeq v0.6.1p1 (20) at default settings. For each sample, per gene read counts were normalized using the default DESeq2 settings (21). Genes showing normalized count values greater than 4 in at least 8 out of 10 populations, within at least one of the treatment types, were kept and the rest were discarded. With these normalized gene count values, we used the linear mixed effects model featured in Barter et al. (12) to determine which genes were differentially expressed between our two selection regimes while accounting for any block effects that may be associated with different rounds of extraction and sequencing (22). Statistical significance for differential expression of any given gene was set at a 5% false discovery rate (FDR) for ~4000 tests, *i.e*. the number of expressed genes that passed filtering (23). The normalized gene count values for the differentially expressed genes were then used as inputs in our analyses.

### Phenotypic Data

We relied on age specific adult mortality and fecundity data for the ten A and ten C populations (13). Mortality and fecundity data were available over the entire adult lifespan of the flies. In our analyses, we focused on average fecundity and logged transformed mortality measures taken over 3-day intervals.

### FLAM Analyses

To find gene regions that might be causally related to the phenotypes studied, we used a statistical learning method called the “fused lasso additive model” or “FLAM” (10). This method has been shown to effectively identify causal loci of phenotypic variation in experimentally evolved populations that exhibit large phenotypic differences (11). In addition, FLAM has the ability to distinguish between these causal loci and those that show genetic differentiation between populations but are *not* causally related to the phenotype of interest (11). We also implemented a permutation procedure for expanding the list of causative loci (11), In this study, a total of 100 permutations of the columns of genetic data were done and the final list consisted of genetic variants which occurred at a frequency of at least 50% among these lists of causative loci.

## Author Contributions

T.T.B. conducted all the analyses presented in this manuscript, was chiefly responsible for the writing of the manuscript, and produced all the figures found in the manuscript.

Z.S.G. contributed by helping with the realignment of the SNP data and was chiefly responsible for altering the SNP data for the FLAM analyses.

M.A.P. conducted the realignment of the SNP data and the statistical tests on the newly aligned SNP data.

J.M.R. contributed by providing critical input on the details of the *Drosophila* transcriptome and assisted in interpreting the genomic causal sites for transcriptomic differentiation.

M.R.R. contributed by constructing the idea of the project and introducing which comparative analyses should be done. M.R.R. also contributed in the writing of the manuscript.

L.D.M. contributed by testing and refining the FLAM analyses for detecting causal molecular regions.

