## Supplementary Materials for "Genome-Wide Architecture of Adaptation in Experimentally Evolved *Drosophila*"

**Supplementary Figures**

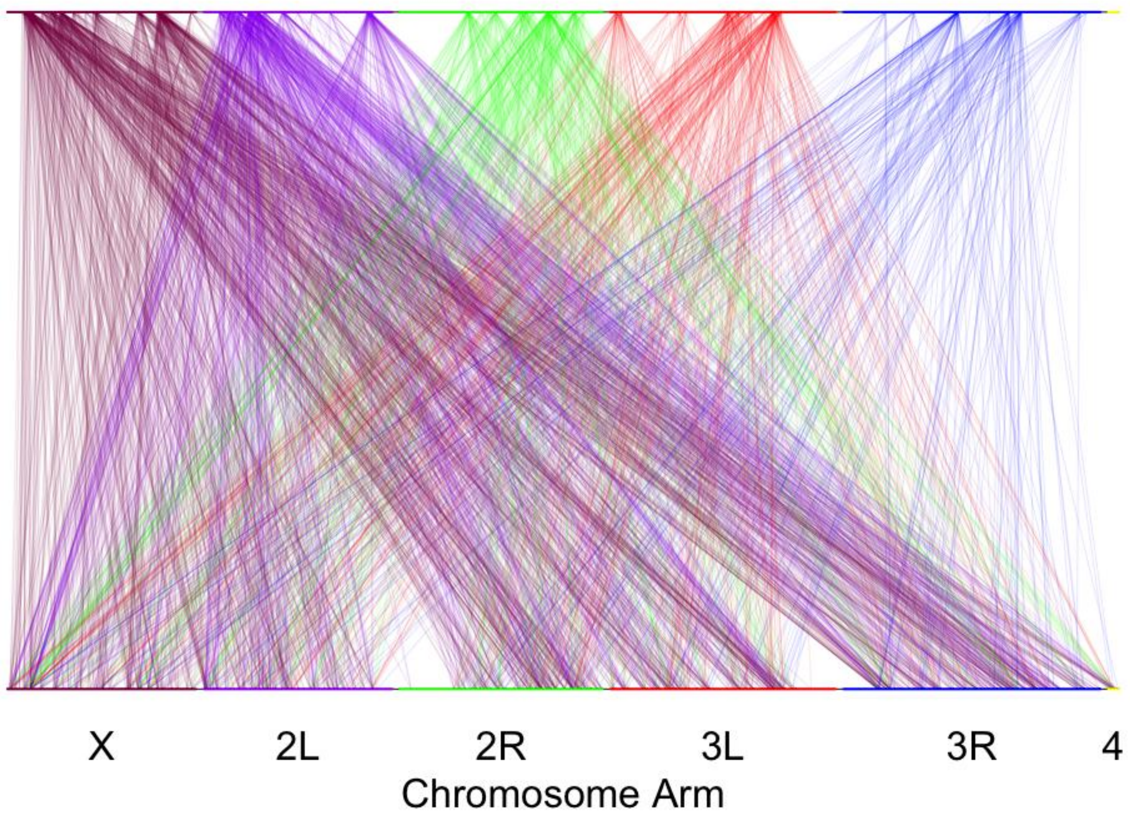


**Fig. S1. Chromosomal distribution of the causal interactions between predictive SNP regions and differentially expressed genes at day 14.** Each line is an interaction between a predictive SNP region (top) to a gene (bottom) for which can accurately predict its differential expression. The color of the line denotes which chromosome arm the genomic region originate from. The genomic regions are classified as 50kb windows in which there contained at least 3 differentiated SNPs between the populations types A and C. The transcripts are classified as the expression of each differentially expressed gene.

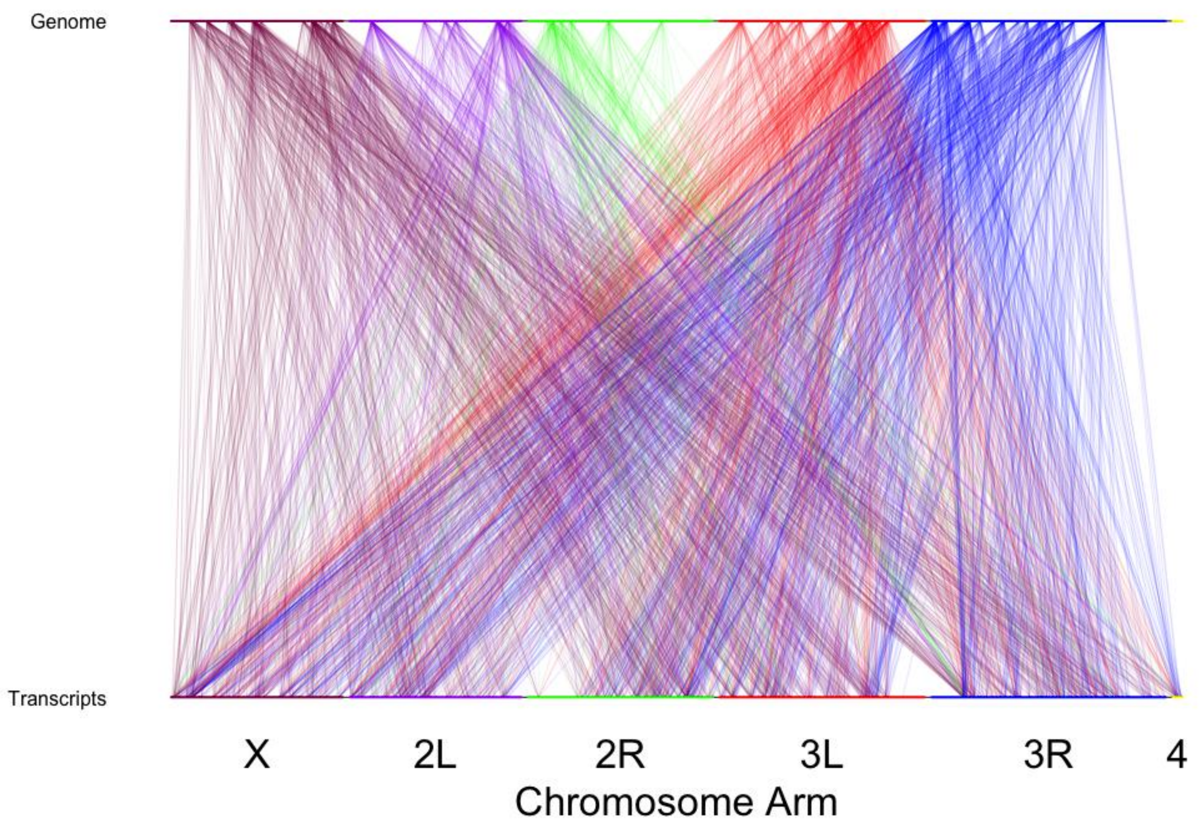


**Fig. S2. Chromosomal distribution of the causal interactions between predictive TE abundance and differentiated transcripts at day 14.** Each line is an interaction between a predictive TE abundance to a gene that the TE abundance can accurately predict the expression. The color of the line denotes which chromosome arm the TE originates from. The TEs are classified as the frequency in which the known TE was found in the population. The transcripts are classified as the expression of each differentially expressed gene.

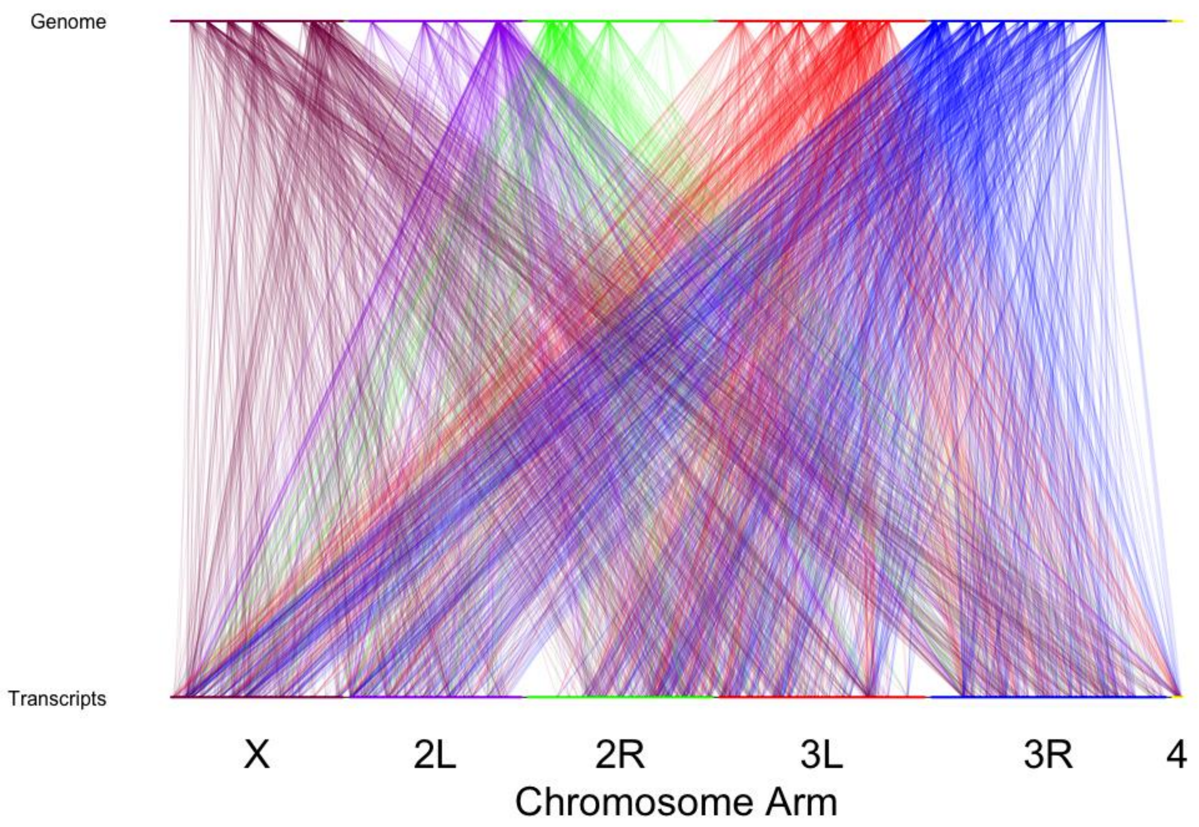


**Fig. S3. Chromosomal distribution of the causal interactions between predictive TE abundance and differentiated transcripts at day 21.** Each line is an interaction between a predictive TE abundance to a gene that the TE abundance can accurately predict the expression. The color of the line denotes which chromosome arm the TE originates from. The TEs are classified as the frequency in which the known TE was found in the population. The transcripts are classified as the expression of each differentially expressed gene.

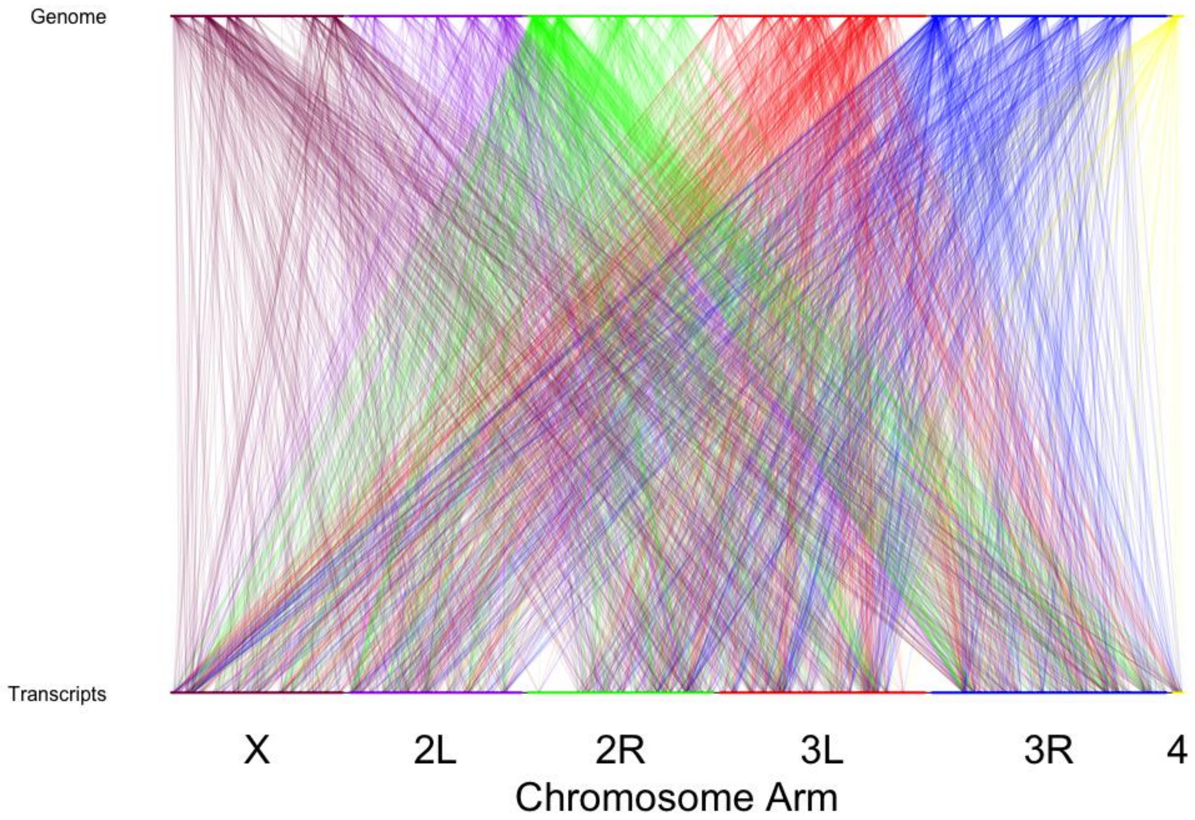


**Fig. S4. Chromosomal distribution of the causal interactions between predictive insertions/deletions and differentiated transcripts at day 14.** Each line is an interaction between a predictive insertion/deletion to a gene that the frequency of the insertion/deletion can accurately predict the expression. The color of the line denotes which chromosome arm the insertion/deletion originates from. The insertions/deletions are classified as the frequency in which the insertions/deletions were found in the population. The transcripts are classified as the expression of each differentially expressed gene.

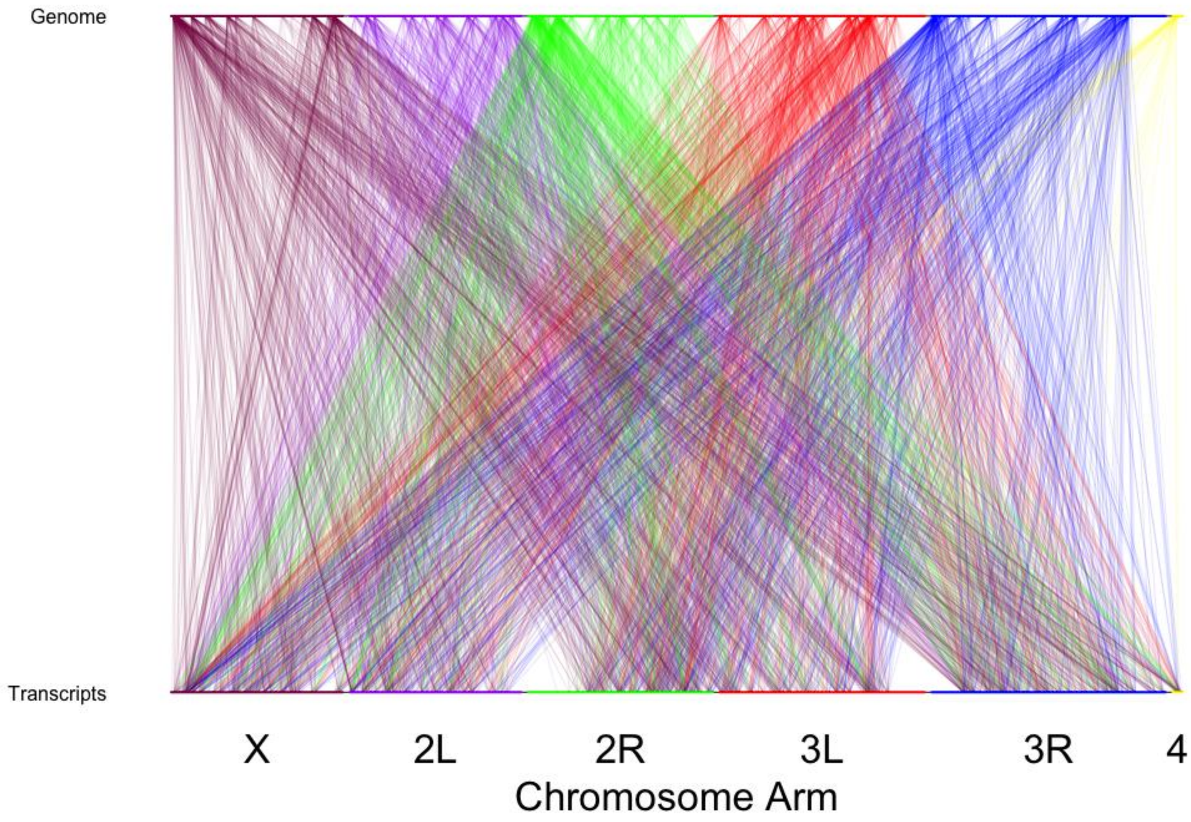


**Fig. S5. Chromosomal distribution of the causal interactions between predictive insertions/deletions and differentiated transcripts at day 21.** Each line is an interaction between a predictive insertion/deletion to a gene that the frequency of the insertion/deletion can accurately predict the expression. The color of the line denotes which chromosome arm the insertion/deletion originates from. The insertions/deletions are classified as the frequency in which the insertions/deletions were found in the population. The transcripts are classified as the expression of each differentially expressed gene.

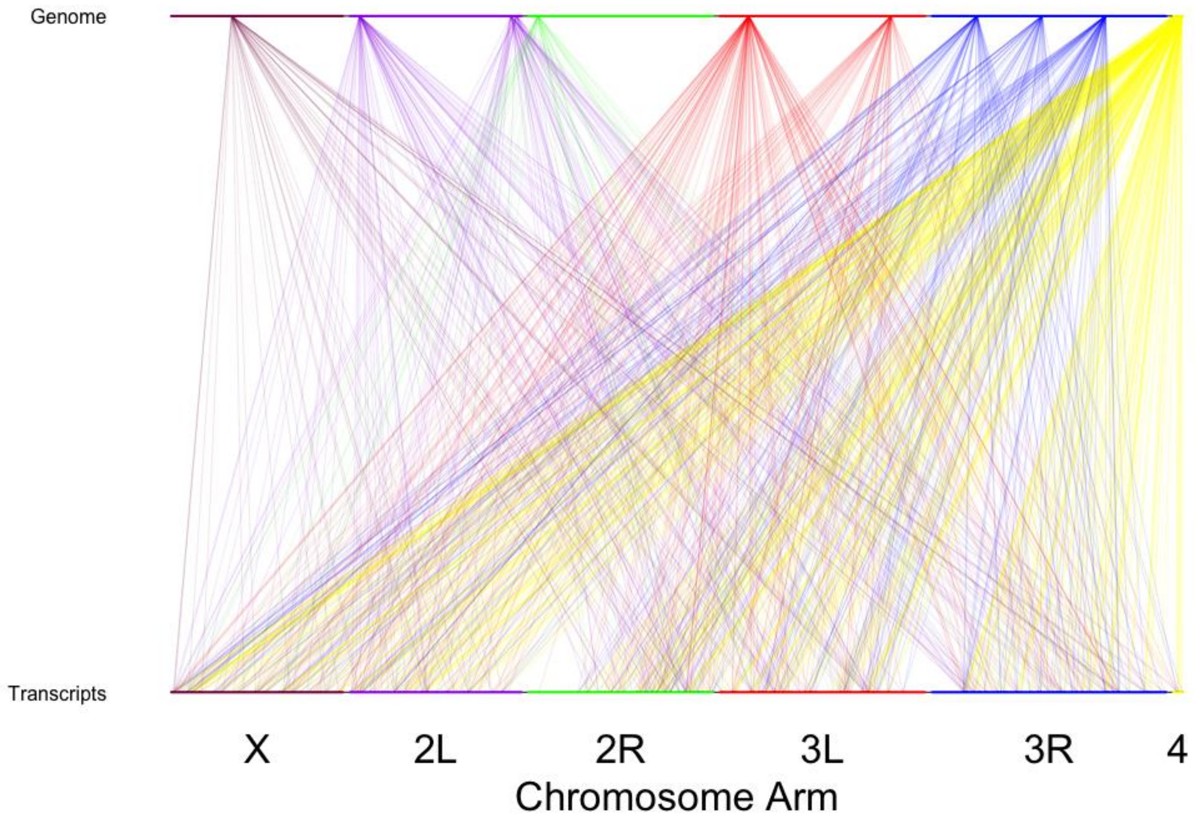


**Fig. S6. Chromosomal distribution of the causal interactions between predictive duplications and differentiated transcripts at day 14.** Each line is an interaction between a predictive duplication to a gene that the frequency of the duplication can accurately predict the expression. The color of the line denotes which chromosome arm the duplication originates from. The duplications are classified as the frequency in which the duplications were found in the population. The transcripts are classified as the expression of each differentially expressed gene.

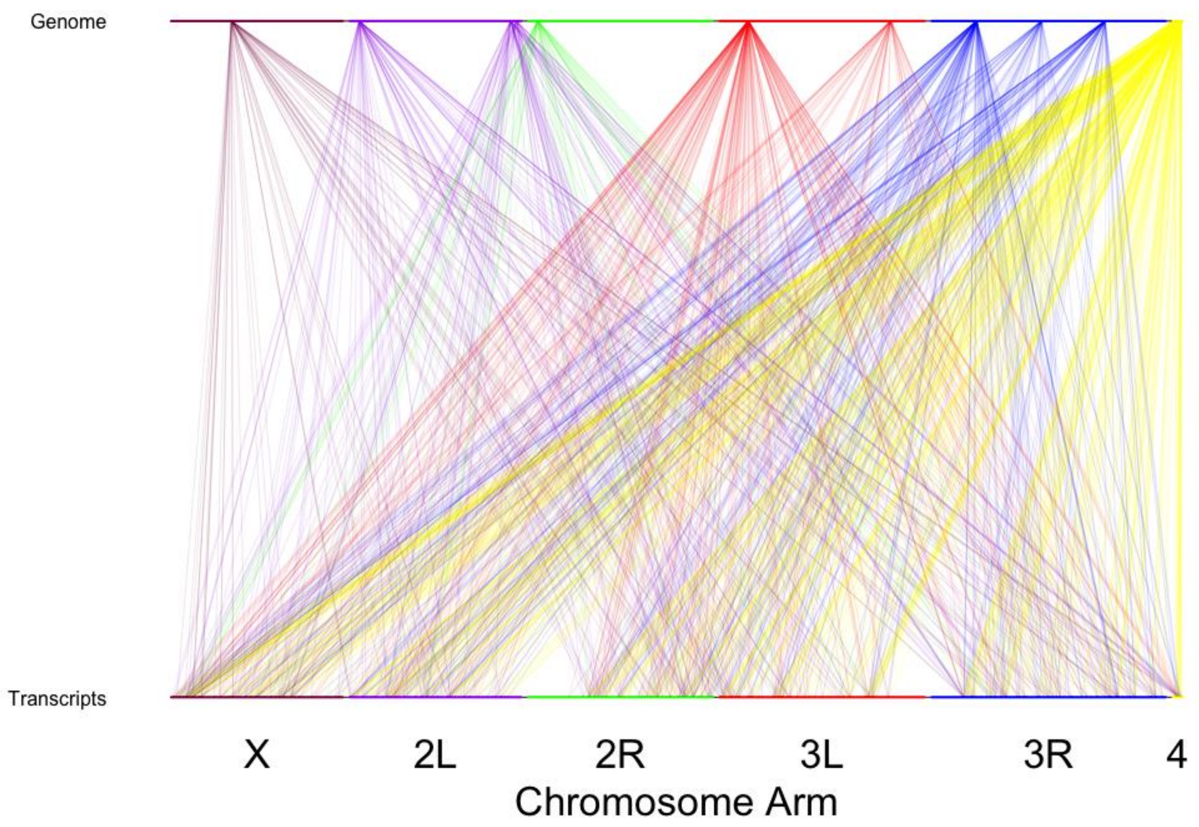


**Fig. S7. Chromosomal distribution of the causal interactions between predictive duplications and differentiated transcripts at day 21.** Each line is an interaction between a predictive duplication to a gene that the frequency of the duplication can accurately predict the expression. The color of the line denotes which chromosome arm the duplication originates from. The duplications are classified as the frequency in which the duplications were found in the population. The transcripts are classified as the expression of each differentially expressed gene.

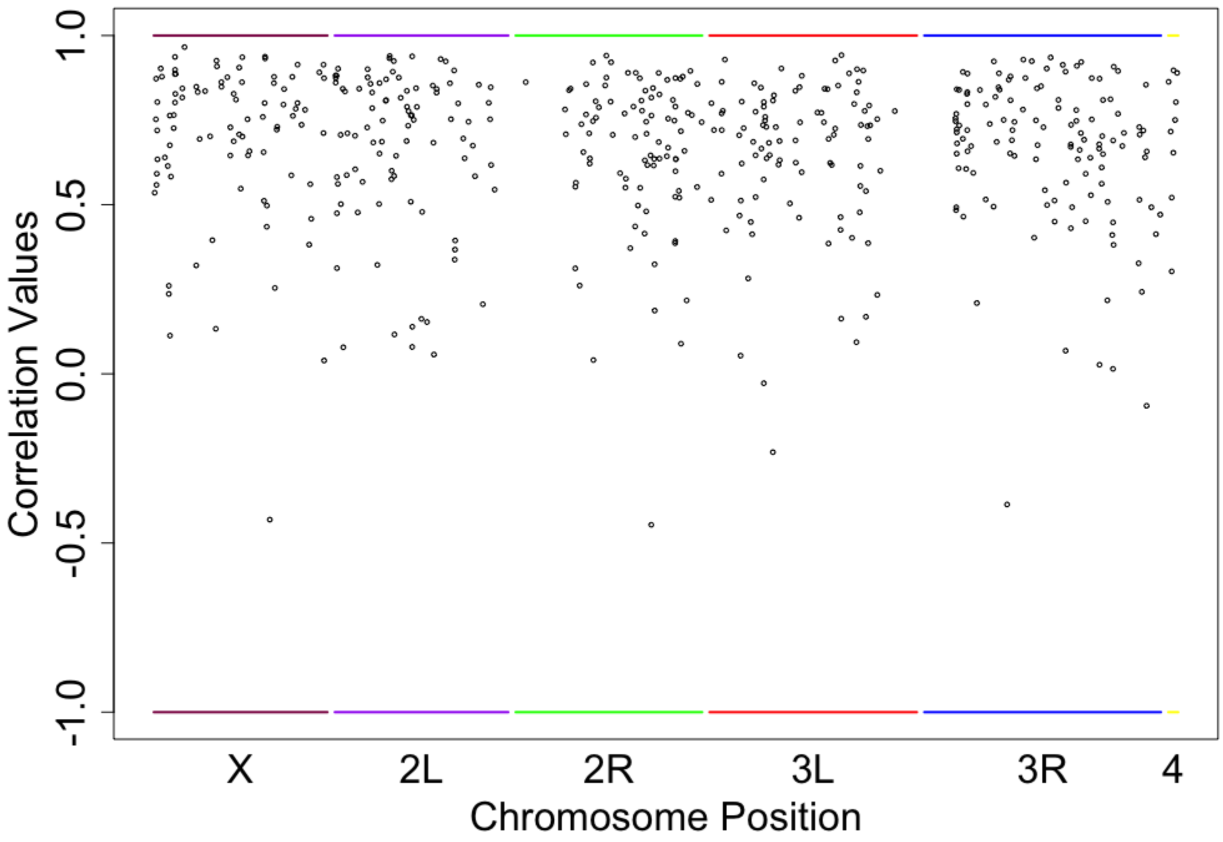


**Fig. S8. Correlation between predicted value and actual value for each differentially expressed gene.** Correlation values were calculated using the predicted expression of each differentiated gene from SNP frequencies and the actual expression values of each differentiated gene at day 14. Each correlation value was plotted at the location of the differentiated gene above.

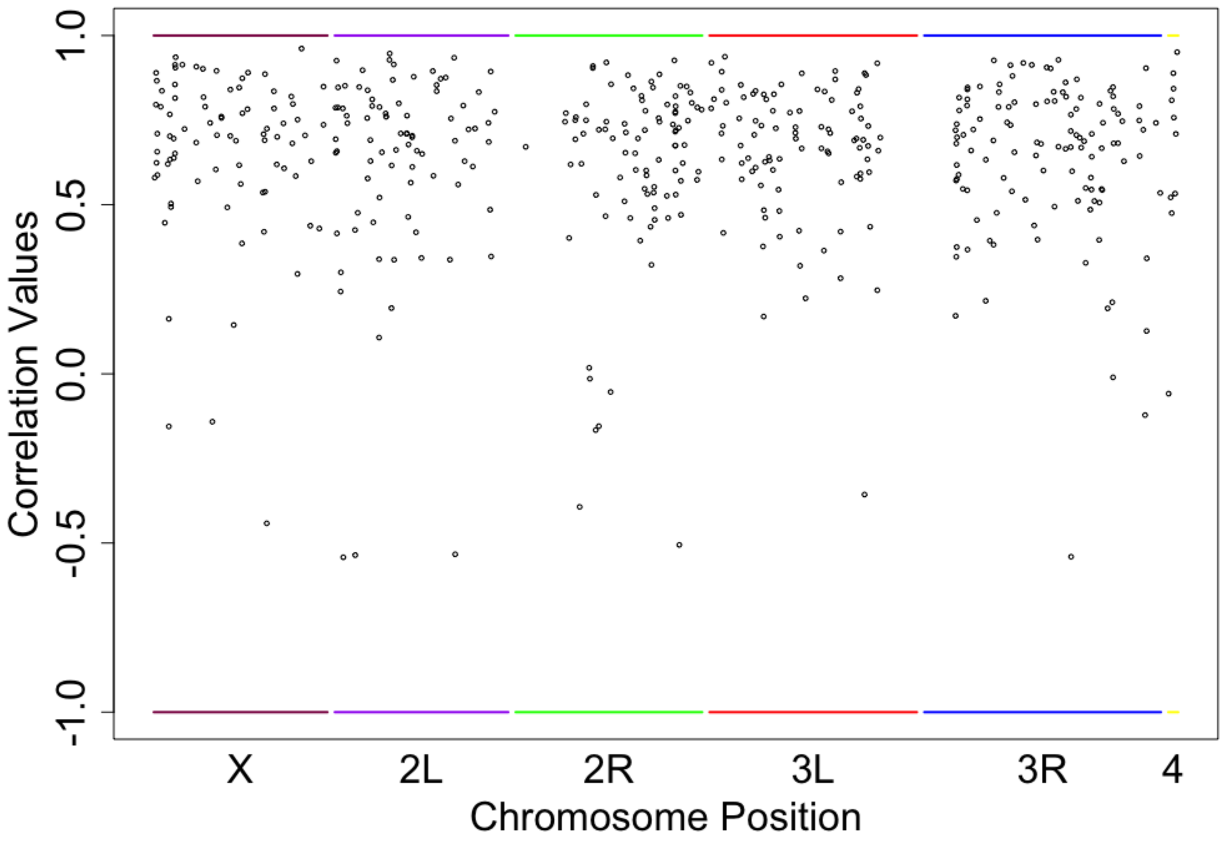


**Fig. S9. Correlation between predicted value and actual value for each differentially expressed gene.** Correlation values were calculated using the predicted expression of each differentiated gene from TE abundance frequencies and the actual expression values of each differentiated gene at day 14. Each correlation value was plotted at the location of the differentiated gene above.

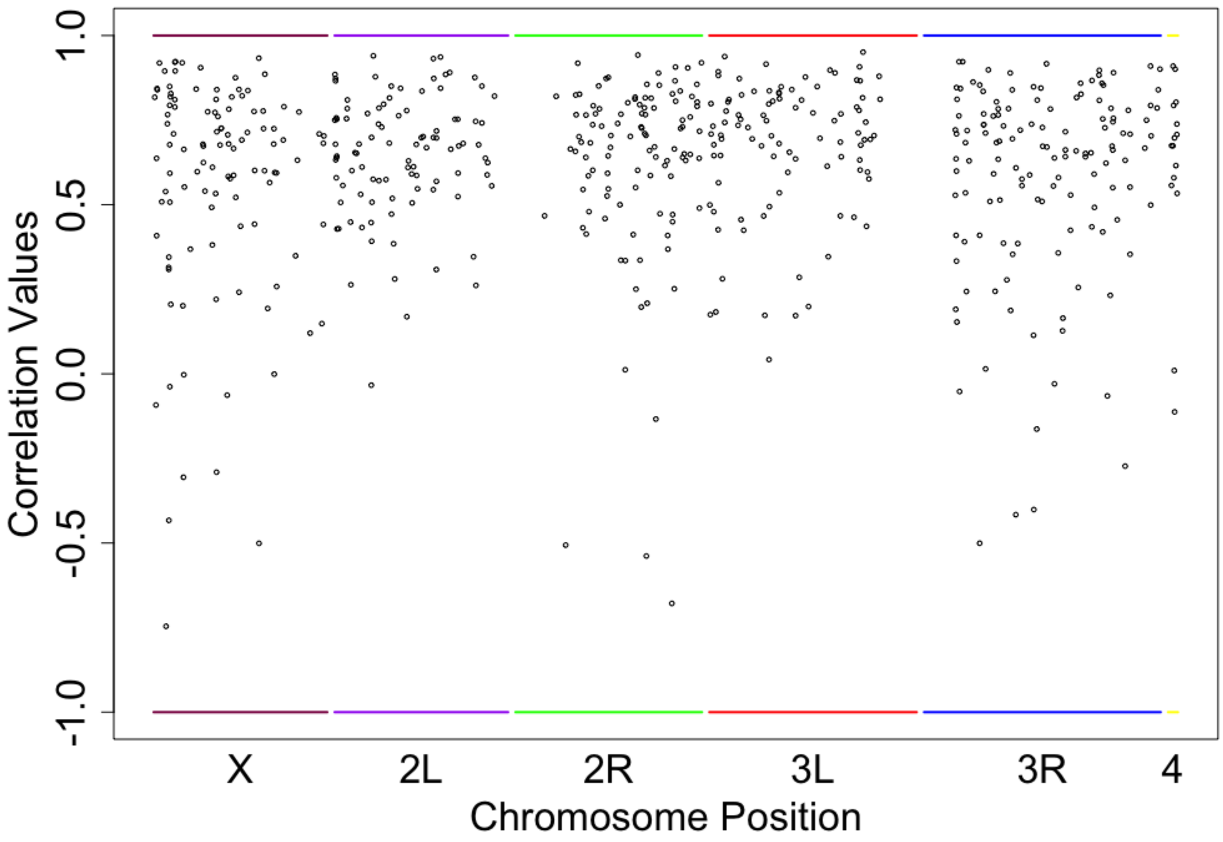


**Fig. S10. Correlation between predicted value and actual value for each differentially expressed gene.** Correlation values were calculated using the predicted expression of each differentiated gene from TE abundance frequencies and the actual expression values of each differentiated gene at day 21. Each correlation value was plotted at the location of the differentiated gene above.

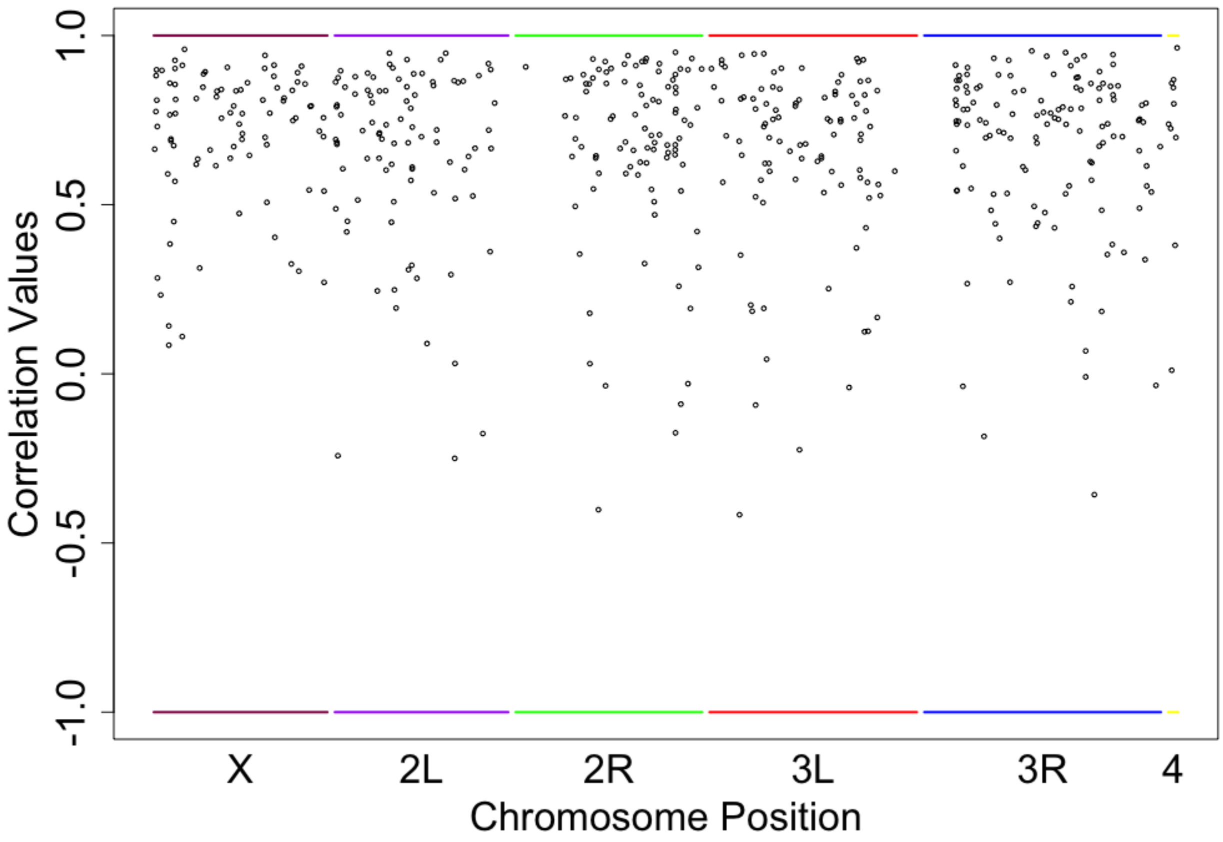


**Fig. S11. Correlation between predicted value and actual value for each differentially expressed gene.** Correlation values were calculated using the predicted expression of each differentiated gene from insertion/deletion frequencies and the actual expression values of each differentiated gene at day 14. Each correlation value was plotted at the location of the differentiated gene above.

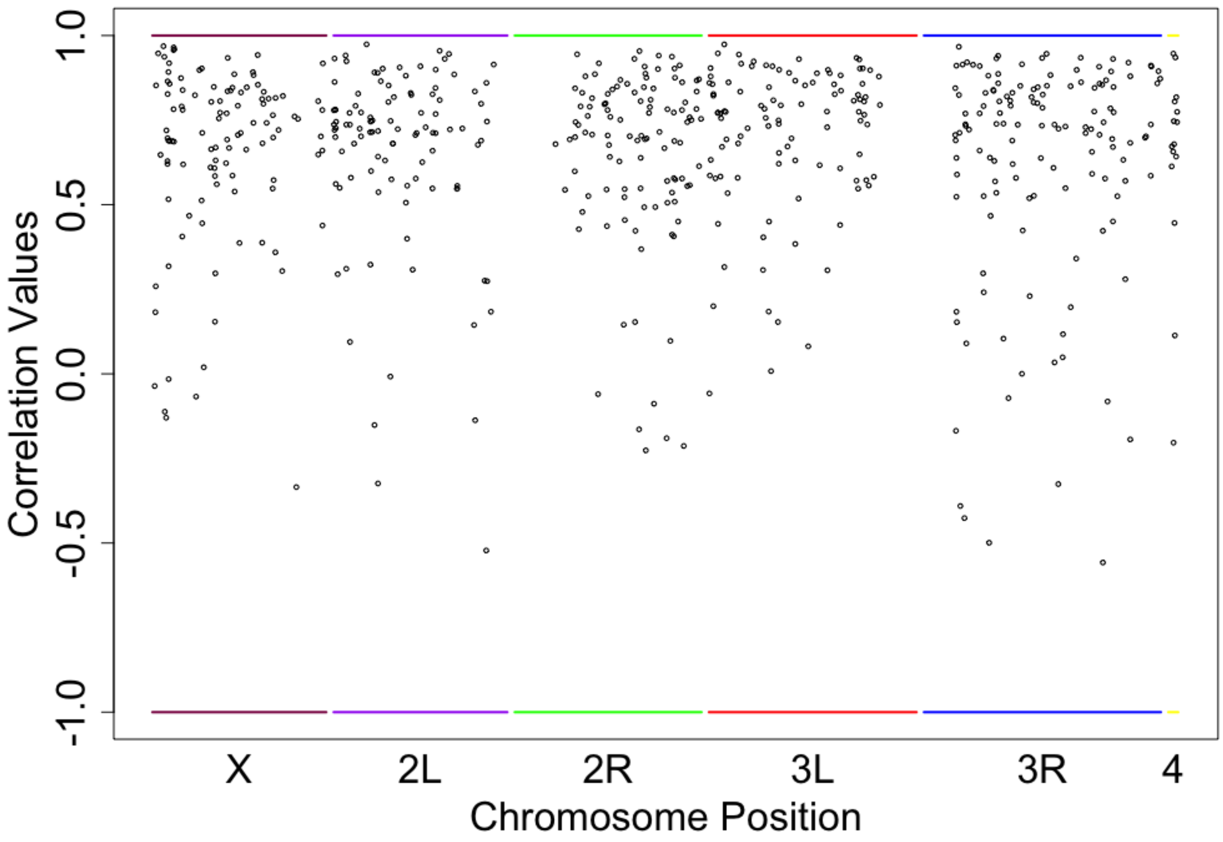


**Fig. S12. Correlation between predicted value and actual value for each differentially expressed gene.** Correlation values were calculated using the predicted expression of each differentiated gene from insertion/deletion frequencies and the actual expression values of each differentiated gene at day 21. Each correlation value was plotted at the location of the differentiated gene above.

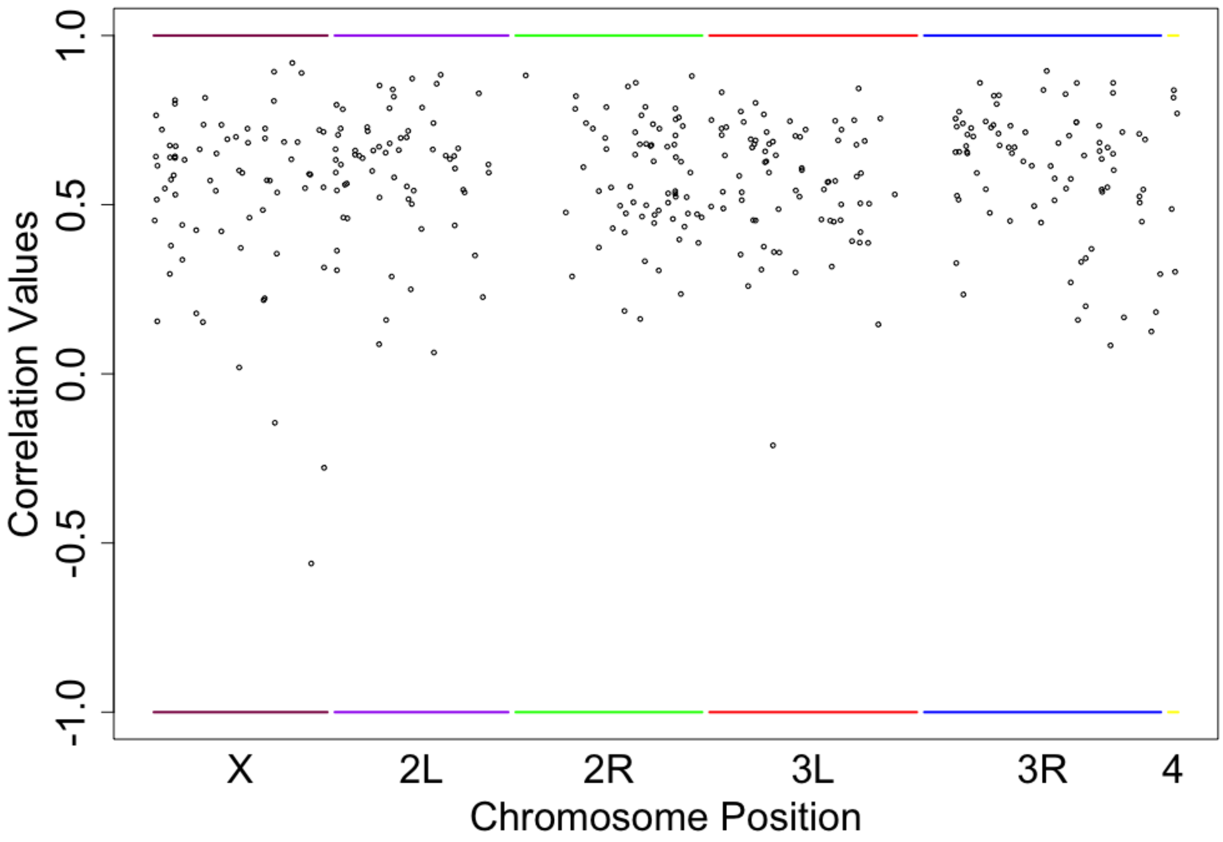


**Fig. S13. Correlation between predicted value and actual value for each differentially expressed gene.** Correlation values were calculated using the predicted expression of each differentiated gene from duplication frequencies and the actual expression values of each differentiated gene at day 14. Each correlation value was plotted at the location of the differentiated gene above.

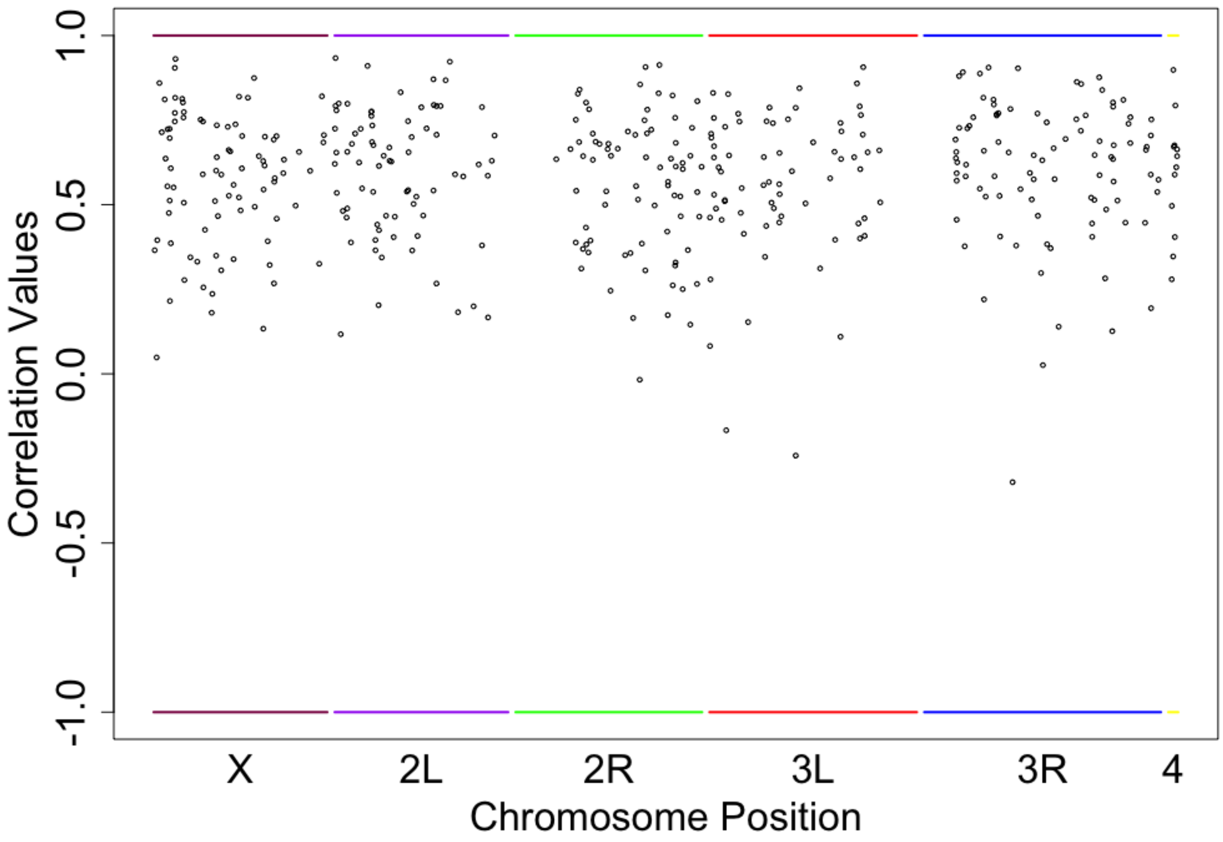


**Fig. S14. Correlation between predicted value and actual value for each differentially expressed gene.** Correlation values were calculated using the predicted expression of each differentiated gene from duplication frequencies and the actual expression values of each differentiated gene at day 21. Each correlation value was plotted at the location of the differentiated gene above.

| Genomic Feature | Day 14 | Day 21 |
|---|---|---|
| **SNP** | 5.9 | 5.7 |
| **TE Insertions** | 5.6 | 6.4 |
| **Small Indels** | 6.6 | 6.6 |
| **DNA Duplications** | 2.7 | 2.8 |

**Table S1. The average number of predictors for each differentiated gene using different genomic features for both day 14 and day 21.**
